# Alternative polyadenylation in the brain is altered by chronic ethanol exposure in a sex- and cell type-specific manner

**DOI:** 10.64898/2026.03.17.712352

**Authors:** Petar N. Grozdanov, Laura B. Ferguson, Brent R. Kisby, Clinton C. MacDonald, Robert O. Messing, Igor Ponomarev

**Author notes:** To whom correspondence should be addressed: Petar N. Grozdanov Texas Tech University Health Sciences Center, 3601 4^th^ Street Lubbock, TX 79430, USA.

## Abstract

Alternative polyadenylation (APA) is a common posttranscriptional mechanism to regulate gene expression. APA generates mRNAs with varying lengths of 3′ UTRs or transcripts that encode distinct protein carboxy-terminal ends. APA is especially important in neurons, where different mRNA variants are often asymmetrically localized to dendrites and axons, and can be locally translated into proteins. Local protein synthesis is crucial for axon guidance, synaptic plasticity, and learning and memory, key processes associated with the development of alcohol use disorder (AUD). We investigated the role of APA in AUD using a mouse model of alcohol dependence characterized by increased voluntary drinking after chronic intermittent ethanol (CIE) exposure. We examined APA during protracted withdrawal from alcohol in three brain regions of male and female mice. Our analyses revealed hundreds of genes undergoing APA in males, but substantially fewer in females, suggesting sex-specific effects of CIE on APA. Notably, male and female mice displayed distinct APA signatures. APA genes were different from differentially expressed genes (DEGs), suggesting that these molecular processes are regulated independently. We also determined that the expression of APA genes was associated with neurons, while DEGs were associated with non-neuronal cells. Many of the APA genes were involved in synaptic integrity, neuroplasticity, and neuronal maintenance, which was consistent with their enrichment in neurons. Our study suggests that APA is a crucial sex- and cell type-specific mechanism in AUD with the potential to influence localized neuronal protein expression during protracted withdrawal and to modify alcohol consumption behavior.

**HIGHLIGHTS:**

- Chronic ethanol exposure in mice results in profound changes of APA genes in brain.
- Commonly regulated cleavage and polyadenylation sites and genes were identified in male but not in female mice.
- There was a minimal overlap between APA and differentially expressed genes (DEGs).
- APA genes were primarily associated with neurons, whereas DEGs were associated with non-neuronal cells.

## INTRODUCTION

Diversity of the eukaryotic transcriptome is attained through a series of alternative messenger RNA (mRNA) processing events leading to the formation of mature mRNAs: alternative transcription 5′ start sites, 3′ end processing, and alternative exon splicing (*1-3*). Alternative 3′ end processing (alternative polyadenylation, APA) is a widespread mechanism that regulates gene expression during normal physiological development and disease progression (*4*). APA of eukaryotic mRNAs involves co-transcriptional endonucleolytic cleavage at specific sites on nascent mRNAs followed by addition of a polyadenosine [poly(A)] tail to the upstream cleaved product—cleavage and polyadenylation, or CPA (*5, 6*). APA can result in three possible outcomes: *(i)* it can produce mRNAs with alternative last exons, thus regulating the size and composition of the proteins by altering the carboxy-terminal coding sequences [also known as intronic APA (IPA)]; *(ii)* in some cases APA can produce non-coding RNAs (ncRNAs) that often have uncharacterized functions; or *(iii)* APA can alter the composition of 3′ untranslated regions (UTRs), which control the stability and localization of mRNAs within cells by inclusion or exclusion of *cis*-regulatory sequences (*4, 7*). Finally, the poly(A) tail ensures stability of the mature transcripts and efficient translation (*2*).

The diversity and abundance of gene transcripts generated by APA and their cognate protein isoforms is highest in the brain, specifically within neurons. APA contributes to dendritic and synaptic development, long-term potentiation, and synaptic plasticity (*7-9*). Neurons address the requirements of protein synthesis in response to external stimuli by transporting mRNAs and translation machinery to synapses, which enables localized protein production (*10*). As a result, neurons possess the synaptic autonomy to regulate localization, translation, and stability of mRNAs at sites where neuronal signals are generated and transmitted between cells (*10*). In neurons, transcripts with longer 3′ UTRs tend to be transported to neurites and neuropils (*11, 12*), where their translation is regulated by the specific demands of synaptic dynamics and activity (*7, 9*). Conversely, transcripts with shorter 3′ UTRs remain associated with the neuronal bodies. These characteristics of local protein synthesis have been implicated in a variety of brain disorders including alcohol use disorder (AUD) (*13-16*).

AUD is recognized as a chronic disease characterized by an inability of individuals to control and limit excessive alcohol consumption, despite the detrimental effects on their health and socioeconomic status. More than 60% of Americans consume alcohol (ethanol) in different forms and 10% were considered to have AUD (*17, 18*). AUD is often characterized as having major effects on the brain and presents itself with comorbidities, such as major depressive disorder and the development of neuropsychiatric disorders (*19*). Moreover, excessive alcohol consumption promotes other chronic health issues including cancer, cardiovascular disease, and others (*20, 21*). The interplay between AUD and these comorbidities underscores the complex nature of alcohol misuse and highlights the need for comprehensive understanding of the physiological changes in the brain occurring during the development of AUD.

Advancements in high-throughput sequencing and bioinformatics have permitted identification of changes in gene expression related to AUD from various brain regions obtained from postmortem human tissues and from animal models of alcohol dependence (*22-24*). However, the full picture of transcriptomic changes in AUD has not been completely captured yet. Here, we used 109 published datasets from a mouse model of alcohol dependence to analyze changes in APA (*25*). In this study, male and female C57BL/6J mice, which have a strong tendency to increase alcohol consumption following development of dependence, were subjected to the chronic intermittent ethanol (CIE) exposure procedure (*25*). The CIE procedure mimics the experience of individuals with AUD, where the subjects undergo episodes of heavy drinking followed by withdrawal periods (*26-28*). Samples from male and female C57BL/6J mice subjected to the CIE procedure were harvested from three brain regions and processed for RNA sequencing (*29, 30*). Changes in mRNA abundance were previously determined for these mice after one week of alcohol withdrawal (*25*) when the peak of increased voluntary alcohol intake was the highest (*31*). Our study identified APA as a distinct but complementary mRNA mechanism to regulate gene expression that was sex- and cell type-specific. Our results suggest that APA is an important process in neurons and has the potential to modulate synaptic gene expression in AUD.

## RESULTS

### Identification of moderate to high expressing “on-site” cleavage and polyadenylation (CPA) sites in whole male mouse brains

Previously, we showed that chronic intermittent ethanol exposure (CIE) escalated alcohol consumption in C57BL/6J alcohol-dependent mice (*25*). In the present study, to examine the changes occurring in the brains of these alcohol-dependent mice, we determined the CPA sites expressed in whole male mouse brains using “on-site” RNA-sequencing technology (Fig. 1A and Materials and Methods). The final curated database contained 44,770 CPA sites that had at least three reads in at least three whole brain samples, located in 15,906 genes and non-coding RNAs (ncRNAs). The majority of CPA sites were located in the 3′-most exon (Supplementary Fig. 1 and 2A, Table S1).

**Figure 1.**
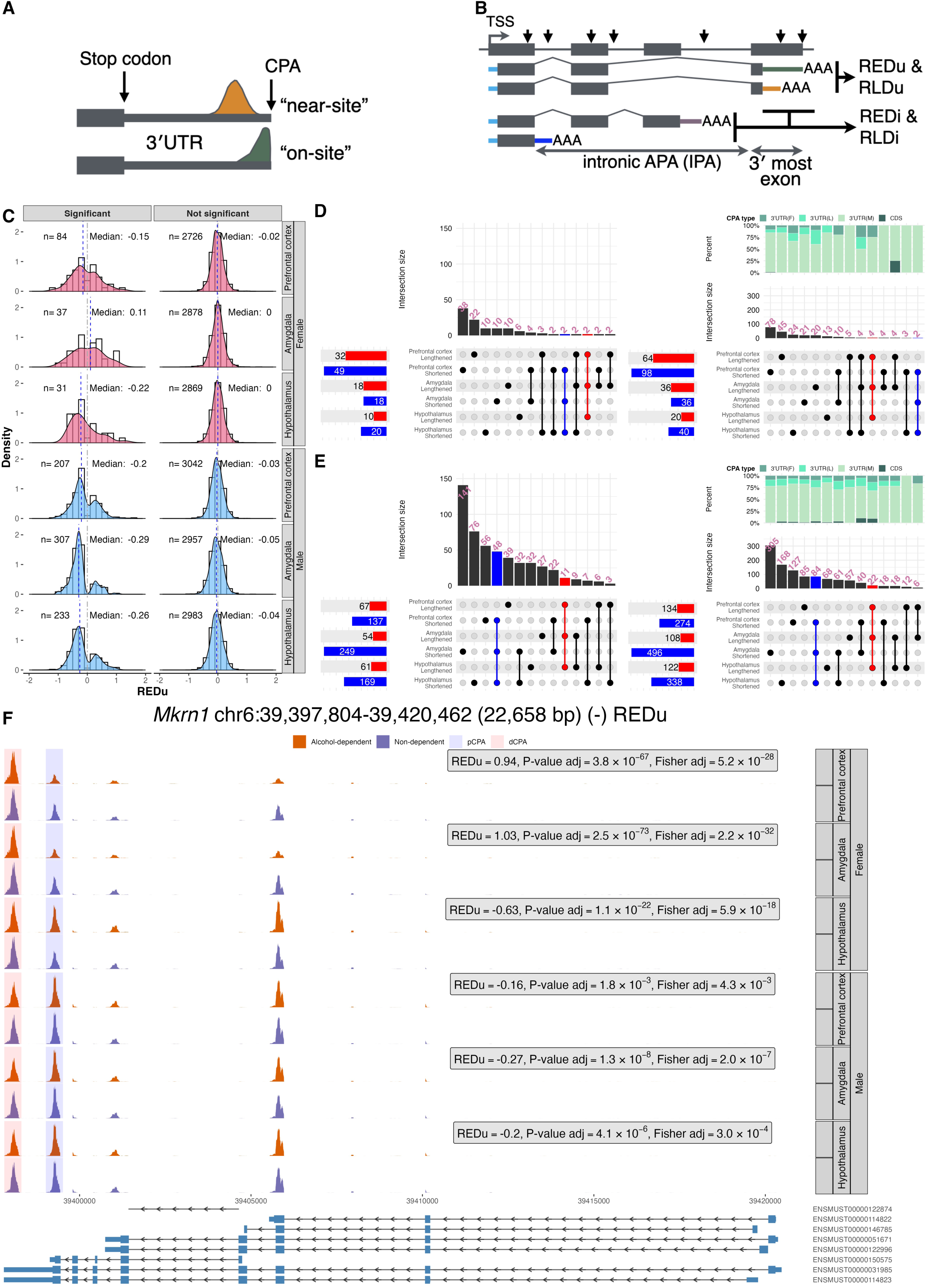
Alternative polyadenylation changes in 3’-most exons (REDu). **A)** Schematic representation of the “near-site” and “on-site” RNA-sequencing technologies. A 3′-most exon containing a single cleavage and polyadenylation (CPA) site is illustrated; stop codon and 3′ untranslated region (3′ UTR) are indicated. The “near-site” read distribution is depicted in orange, while the “on-site” distribution is shown in dark green. **B)** Schematic diagram of CPA site localization, the transcripts produced, and output metrics from the MAAPER bioinformatic tool. Black arrows indicate CPA sites located in various genomic features, such as the 5′-most exon, introns, internal exons, single exons, and 3′-most exons. REDu and REDi represent the log_2_ relative expression difference scores for APA events in the 3′-most exon (REDu) and intronic (REDi) sites (including introns, internal exons, etc.). Transcription Start Site (TSS) is indicated. **C)** Density distributions are presented with histogram plots of genes exhibiting statistically significant REDu (left) and non-significant (right) scores for the indicated brain region and sex. **D)** UpSet plots illustrate unique and overlapping genes (left) and CPA sites, the two with biggest change of ratio (right) across brain regions for shortened and lengthened APA genes in female mice with statistically significant REDu scores. Overlapping genes that lengthened across all brain regions are highlighted in red, while shortened genes are represented in blue. The percentage of CPA types (various shades of green) is indicated for each overlap. **E)** The same representation as in **D)** but for male mice, showing an increased number of overlapping genes across all three brain regions. **F)** The *Mkrn1* gene is provided as an example of APA changes in the 3’-most exon. REDu scores, BH-adjusted likelihood ratio and BH-Fisher’s exact tests for each pairwise comparison are presented, with negative REDu scores indicating shortening. Distal CPA (dCPA) and proximal CPA (pCPA) are shown in light blue and red, respectively. Different *Mkrn1* transcripts are displayed along with their Ensembl IDs.

**Figure 2.**
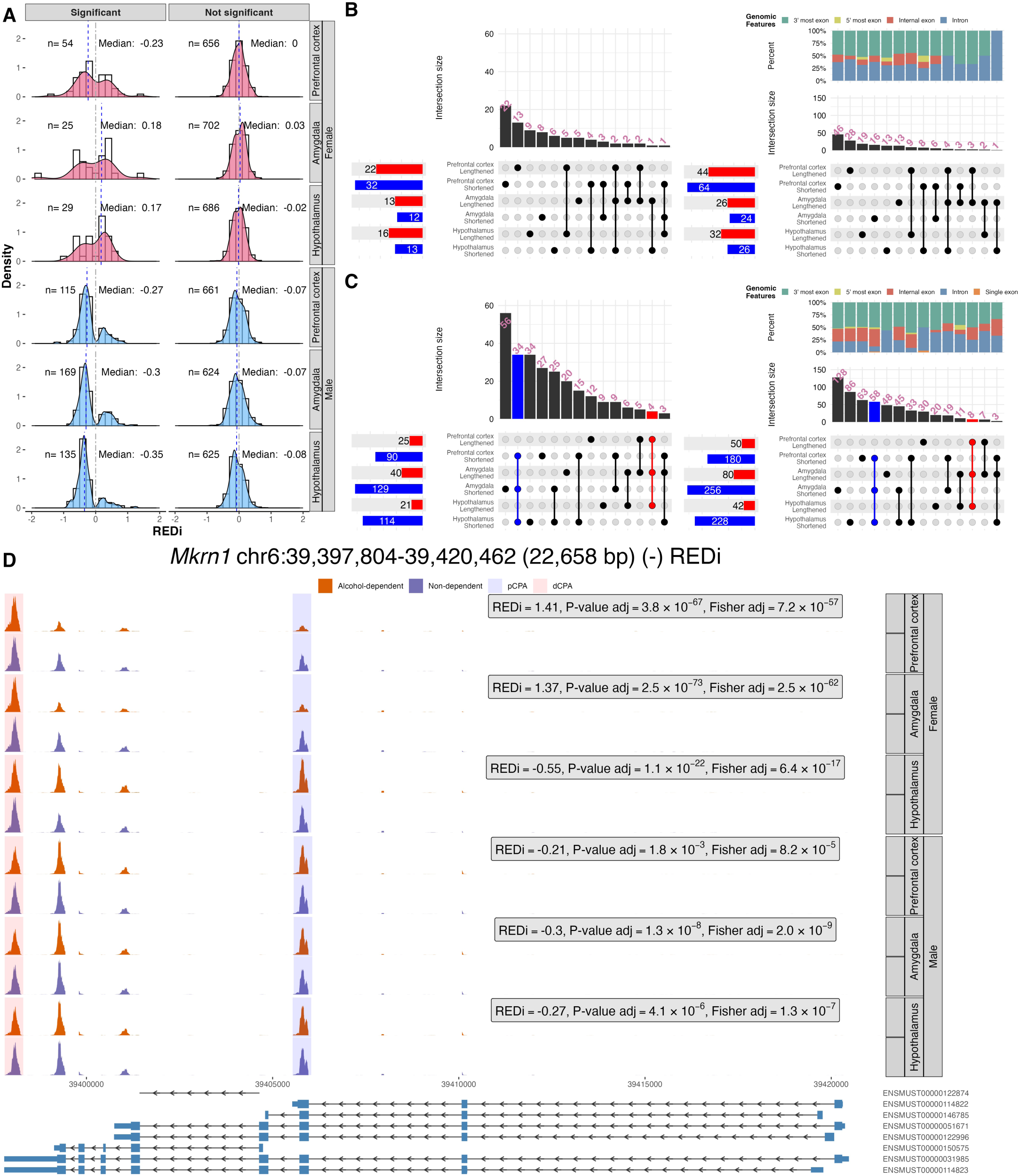
Alternative polyadenylation changes in introns (REDi). **A)** Density distributions and histogram plots are shown for genes with statistically significant REDi (left) and non-significant (right) scores calculated for the indicated brain region and sex. **B)** UpSet plots depict unique and overlapping genes (left) and CPA sites (right) across brain regions for shortened and lengthened APA genes in female mice with statistically significant REDi scores. Overlapping genes that lengthened across all brain regions are shown in red, and shortened genes are in blue. The percentage of genomic feature events is indicated for each overlap. **C)** The same representation as in **B)** but for male mice, again note the increased number of overlapping genes in all three brain regions. **D)** The *Mkrn1* gene serves as an example of intronic APA changes. REDi scores for each pairwise comparison is presented, with negative REDi scores indicating shortening and positive scores indicating lengthening. Distal CPA (dCPA) and proximal CPA (pCPA) are represented in light blue and red, respectively. Different *Mkrn1* transcripts are displayed along with their Ensembl IDs.

### TagSeq is a reliable RNA sequencing technology for APA identification

We proceeded to determine differential CPA site usage (alternative polyadenylation, APA) in alcohol-dependent relative to non-dependent mice, using 109 samples obtained from three critical brain areas important for the development of alcohol dependence: prefrontal cortex, the amygdala, and hypothalamus (*32*). These datasets were generated using TagSeq, an RNA-Seq method that prioritizes mRNA 3′ ends (*29, 30*) resulting in “near-site” RNA-sequencing reads (Fig. 1A, Supplementary Fig. 1C). Although TagSeq focuses on 3′ ends using a “near-site” approach, it has not yet been shown that it can be reliably used for APA analysis. Using orthogonal bioinformatic approaches (e.g., MAAPER, and metagene plots), we showed that the relative normalized distribution of the reads in TagSeq versus the “on-site” reads, the distribution across CPA types, the number of training genes, and the uniformity of reads distribution were consistent across brain regions, sexes, and experimental groups (Supplementary Fig. 1 and 2). Thus, our analyses revealed that TagSeq could be reliably used to examine APA changes in alcohol dependence and could likely be applied in other experimental designs.

### APA changes profoundly in male but not female mice in an alcohol-dependence model

MAAPER examines two types of APA, APA located in the 3′-most exon and APA located in intronic regions (IPA); the latter includes APA events located in introns, internal exons and 5′-most exons (Fig. 1B) (*33*). MAAPER produces relative expression difference (REDu) and relative length difference (RLDu) scores for APA in the 3′-most exon sites, and REDi and RLDi scores for IPA sites (Table S2, Fig. 1 and 2). In our APA and IPA analyses, we considered changes to APA in a gene to be statistically significant when both Benjamini-Hochberg’s-corrected likelihood ratio and Fisher’s exact tests were smaller than or equal to 0.05 (for details see Materials and Methods).

The majority of the CPA sites were localized in the 3′-most exons, followed by the introns across all pairwise comparisons. The distribution of CPA sites across the genomic features [5′-most exon (0.2%), intron (5.6%), internal exon (2.0%), single exon (2.5%), and 3′-most exon (89.6%)] on average was similar in all samples (Supplementary Fig. 2A). This distribution closely resembled the observed distribution in the curated brain polyA mm10 database, albeit on average the intronic sites were less represented in the comparisons (17.5% vs. 5.6%, respectively, Supplementary Fig. 2A). The numbers of predicted CPA sites per gene were also similar across the comparisons, with more than half of the genes having a single CPA site (Supplementary Fig. 2E).

Next, we plotted density graphs for each pairwise APA and IPA comparison across brain region, sex, and statistical significance. Male mice in every brain region showed overall shortening for the genes changing CPA sites in the 3′-most exons (Fig. 1C) and for the IPA genes (Fig. 2A). The amygdala was the brain region that showed the most genes and ncRNAs that changed APA in the 3′-most exon (307 events) and IPA (169), followed by hypothalamus (233 and 135) and prefrontal cortex (207 and 115 events respectively, Fig. 1C). Negative median REDu scores, which describe the relative differences in expression between the two most differentially expressed transcripts within the 3′-most exon in a gene, imply net shortening of the 3′ UTRs in these gene sets. The median REDu scores were -0.20 for prefrontal cortex, -0.29 for amygdala, and -0.26 for hypothalamus (Fig. 1C). The median REDi scores, which describe the relative change of transcript abundance between a CPA site within the 3′-most exon and either, intron, 5′-most exon, internal exon or intron, were - 0.27 for prefrontal cortex, -0.30 for amygdala, and -0.35 for hypothalamus (Fig. 2A).

A common APA pattern did not emerge in the female mice as it did for male mice (Fig. 1C and 2A). The number of genes changing APA in the 3′-most exons and IPA in female mice was substantially smaller than in male mice. Most genes changing APA and IPA in females were detected in the prefrontal cortex (84 for APA and 54 for IPA), followed by amygdala for APA (37) and hypothalamus for IPA (29). The median REDu scores for APA were -0.15 for prefrontal cortex, 0.11 for amygdala, and -0.22 for hypothalamus, indicating net shortening in prefrontal cortex and hypothalamus but lengthening in amygdala in response to alcohol dependence in female mice.

### APA genes and CPA sites overlap in alcohol-dependent male but not female mice

Next, we wanted to examine whether there was a commonality in the affected APA genes across all three brain regions within the same sex. We separated the statistically significant genes based on the directionality of the change, e.g., shortened or lengthened (REDu and REDi < 0 and >0) across brain region and sex, and performed an overlapping analysis, which was summarized in UpSet plots (*34*) (Fig. 1D, E, 2B and 2C, and Table S3). We excluded ncRNAs and pseudogenes from this analysis, since these RNAs did not have annotated CPA sites in the PolyA_DB (*35*) and ncRNAs/pseudogenes (*1700020I14Rik*, *2210408F21Rik*, *2610005L07Rik*, *Chd3os*, *Malat1*, *Pisd-ps1*, *Tug1*) represented only a small fraction for each comparison. We observed 48 common shortened and 11 common lengthened protein-coding genes in the 3′-most exon category and 34 common shortened and 4 common lengthened IPA genes in male mice. Conversely, female mice showed very limited commonality with only 2 shortened and 2 lengthened in the 3′-most exon and no overlapping IPA genes across the three brain regions.

We asked whether there was an overlap between the CPA sites in 3′-most exons (REDu) and intronic (REDi) analyses. First, we determined the overlap across all CPA sites in all comparisons. Overall, we identified a total of 18,662 unique CPA sites (Supplementary Fig. 3A). From these, 17,349 were present in multiple brain regions and sexes. A relatively small portion (∼7%, 1,313 sites) of the CPA sites were unique per brain region and sex. An even smaller portion of these CPA sites were unique to brain region and sex, and were linked to genes that showed statistical significance based on our conservative threshold (Supplementary Fig. 3B–E). Next, we identified CPA sites of APA genes that were associated with CPA sites identified in all comparisons (Supplementary Fig. 3F and G). These results suggested that these sites had the potential to be regulated in all brain regions but were subject to regulation only in male mice.

We further evaluated the top two CPA sites with biggest change of expression ratio in the statistically significant APA genes. Again, the male mice showed consistent overlap of these CPA sites in both 3′-most exons and IPA genes (Fig. 2B and 2C). We also determined the distribution of CPA sites based on a CPA type. Most overlapping CPA sites in the 3′- most exon were in the 3′UTR(M) CPA type, which designates CPA sites that were not first (F), last (L), or in the CDS (i.e., middle, M). Similarly, CPA sites for the IPA genes showed a common pattern in the male mice. Roughly half of the CPA sites in IPA genes were in the 3′- most exon; the rest were distributed across the 5′-most exons, introns, and internal exons (Supplementary Fig. 2D). A substantial part of the overlapping CPAs in male mice that were located upstream of the 3′-most exon, were found in internal exons. The physiological consequences of this observation were not clear (see Discussion). The female mice did not exhibit common CPA sites that were present in the shortened or lengthened genes in all three brain regions studied (Fig. 2B).

We selected the Makorin ring zinc finger protein 1 (gene symbol: *Mkrn1*) to illustrate the changes in both the 3′-most exon (Fig. 1F) and intronic (IPA, Fig. 2D) events. In addition to its function as an E3 ubiquitin ligase, an alternatively spliced and polyadenylated form of *Mkrn1* has been identified as a poly(A)-binding protein-interacting protein that stimulates translation in neurons (*36*). Overall, the APA regulation of *Mkrn1* proved to be complex. In all pairwise comparisons in males, we observed that the *Mkrn1* transcripts were shortened, illustrated by a transition from the distal CPA (dCPA) to the proximal CPA (pCPA), varying in magnitude based on REDu values (Fig. 1F). In female mice, *Mkrn1* transcripts were lengthened in the prefrontal cortex and amygdala and were shortened in the hypothalamus. The CPA sites that were in the 3′ most exon may have been associated with different transcripts (e.g., ENSMUST00000150575, ENSMUST00000114823 and ENSMUST0000031985). However, we could not confidently determine this association from the current data.

Some of the pCPA sites for *Mkrn1* were annotated to reside within an intron in the curated PolyA_DB (*35*). We were able to link at least one of them to a transcript that produced an alternative open reading frame (ENSMUST00000114822, Fig. 2D). The pCPA in both males and females in all brain regions was at the end of the ENSMUST00000114822 transcript (Fig. 2D, highlighted in blue). However, the dCPA could correspond to either the ENSMUST00000114823 or ENSMUST00000031985 transcript (Fig. 2H, highlighted in red). In the female amygdala and prefrontal cortex, *Mkrn1* underwent a lengthening, as indicated by positive REDi values, while in all other comparisons, it was observed to shorten, albeit to a lesser extent (Fig. 2D).

These results suggest that there were profound APA changes associated with alcohol dependence and withdrawal in male mice in all three brain regions. However, such widespread APA changes were not observed in the female mice. Therefore, we propose that alternative polyadenylation is influenced by alcohol dependence in a sex-dependent manner.

### APA genes are different from DE genes

Next, we examined how many differentially expressed genes (DEGs) determined by the analysis of the original study of *Ferguson et al. 2022* (*25*)] overlapped with APA genes. To have an objective comparison, we mapped the DEGs to the gene symbols that were detected in our APA analyses. This mapping returned fewer DEG gene symbols than in the original study (Supplementary Fig. 4A). DEGs were considered statistically significant based on a nominal p-value ≤ 0.05. The relatively fewer DEGs we observed prompted us to investigate the reasons for this discrepancy from the original study. One possibility was that some DEGs were not mapped because of low abundance, and we excluded them during our APA analysis. To test this hypothesis, we used the average expression from the original study and asked whether the average gene expression was smaller for the genes detected only in the DEG analysis vs the genes detected in both datasets. Indeed, the average gene expression detected in the DEG analysis alone was smaller than the average expression for the genes presented in both analyses (Supplementary Fig. 4A, Wilcoxon’s test p ≤ 0.05). This suggested that our decision to focus on moderate-to-high expressing genes was valid.

We generated proportional Venn diagrams for APA in the 3′-most exons (REDu, Fig. 3A) and IPA (REDi, Fig. 3B) genes relative to DEGs across brain regions and sex. This analysis revealed only a few genes that overlapped between APA in the 3′-most exon or IPA genes and DEGs in male samples from all three brain regions. In females, even fewer genes overlapped only in amygdala and hypothalamus—we noted no overlap between APAs and DEGs for either REDu or REDi in the prefrontal cortex in female mice (Fig. 3A, B). These findings suggest that APA mechanisms regulate gene expression independently of changes in mRNA abundance in alcohol dependence and do so differently in female and male mice.

**Figure 3.**
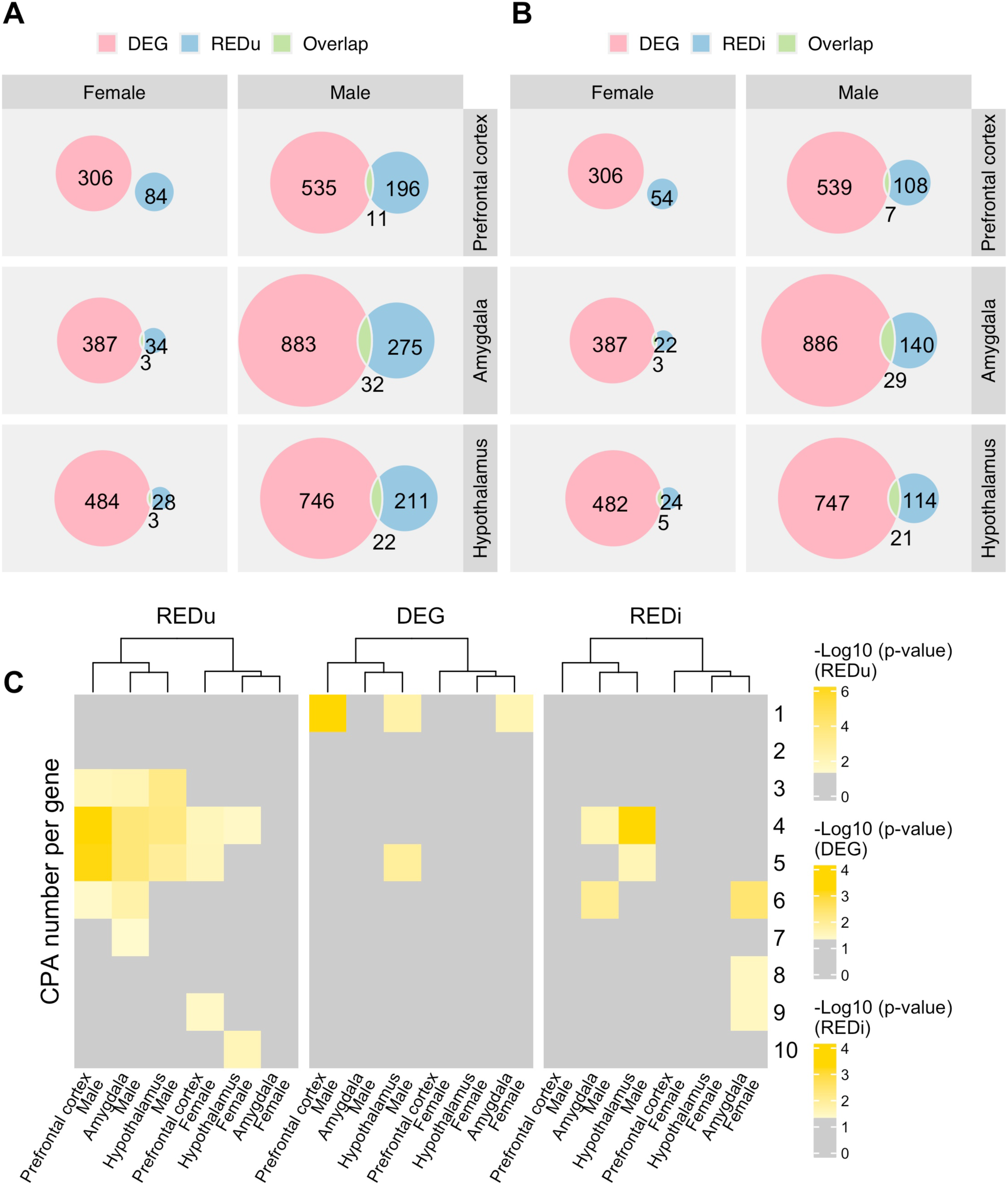
The overlap between APA and DE genes is minimal. **A)** Proportional Venn diagrams illustrate the relationship between APA in the 3’ most exons (REDu) and DE genes that were mapped to the genes detected in MAAPER, analyzed separately for each pairwise comparison. The numbers of unique (blue for APA and salmon for DE) and overlapping (green) genes are shown. **B)** Proportional Venn diagrams depict the relationship between intronic APA (REDi) and DE genes, similarly, mapped to the genes detected in MAAPER for each comparison. The numbers of unique (blue for APA and salmon for DE) and overlapping (green) genes are indicated. **C)** Independent hierarchically clustered heatmaps display hypergeometric probability for enrichment based on the number of CPA sites per gene, considering changes in the 3’-most exonic (REDu), intronic APA (REDi), or DE genes. The color scale corresponding to each hypergeometric distribution is shown on the right. Only the -Log_10_ transformed p-values less than or equal to 0.05 are shown on these color scales.

Next, we wanted to test whether the DEGs were more likely to be associated with genes that have only a single CPA site or genes with multiple CPA sites. To address this question, we used a hypergeometric distribution test to determine enrichment and statistical significance based on number of CPA sites per gene (Fig. 3C). DEGs in male hypothalamus and prefrontal cortex, and female amygdala were significantly enriched (p ≤ 0.05) in the category of single CPA site genes. We performed similar enrichment analyses for the REDu and REDi genes. While most of the significantly enriched APA genes were in genes with three to six CPA sites, the REDu genes in the female prefrontal cortex and hypothalamus were enriched in a category with nine and ten CPA sites, respectively. These results further support the concept that the DEGs and APA genes are regulated by different mechanisms in alcohol dependence.

### APA genes are involved in synaptic integrity, plasticity and maintenance

Gene Ontology (GO) analyses determine the potential roles and functions of regulated genes by comparing them with a comprehensive reference database (*37*). We wanted to investigate the association of the APA and IPA genes across all three categories of GO terms: Biological Process (BP), Cellular Component (CC), and Molecular Function (MF).

The gene lists we examined were categorized by brain region, sex and further divided by directionality based on negative and positive REDu or REDi scores into shortened and lengthened genes. Our analysis revealed 99 unique GO terms (out of a total of 114) for biological processes (BP), 68 (out of 125) for cellular components (CC), and 33 (out of 53) for molecular function (MF) for the REDu gene, and 26 unique GO terms (out of a total of 27) for BP, 37 (out of 52) for CC, and 13 (out of 14) for MF for the REDi genes, all of which were statistically significant (p.adjust ≤ 0.05). Due to the smaller number of APA genes identified in female mice, fewer GO terms were associated with both REDu and REDi genes (Fig. 4A and B, Table S4 and S5). Approximately 78% of the biological process terms associated with shortened APA events occurred in the male hypothalamus, while 52% of the cellular components of the REDi genes were enriched in the male amygdala. As a notable exception, 37% of the biological process terms were enriched for lengthened IPA (REDi) events in the female hypothalamus. (Table S4). However, most of the statistically significant BP and MF terms were more likely to be present in just one of our categories.

**Figure 4.**
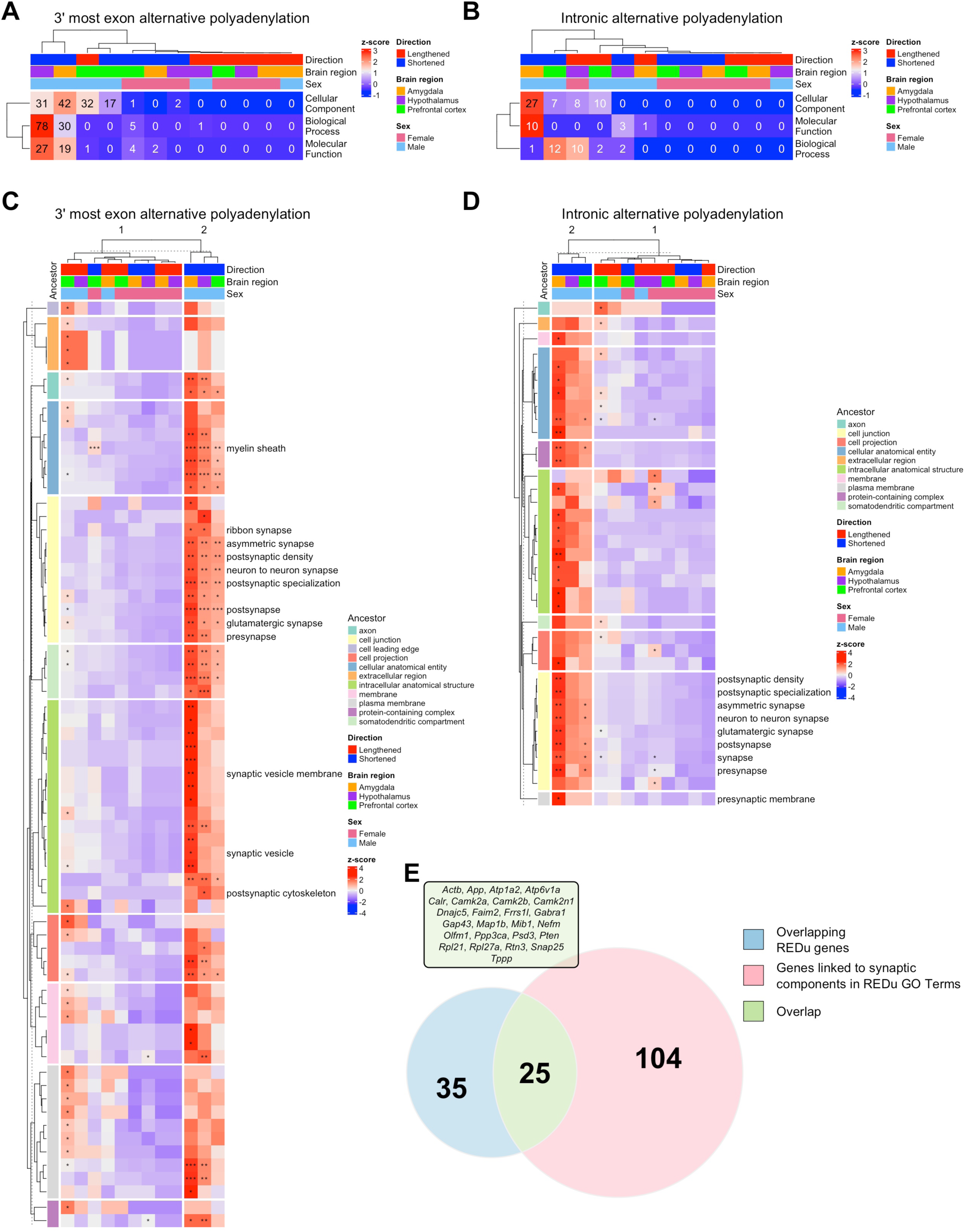
APA genes in the 3′-most exons and introns are functionally enriched for synaptic cellular components. **A)** Z-score normalized hierarchically clustered heatmap summarizing the number of statistically enriched (p.adjust ≤ 0.05) Gene Ontology (GO) terms in at least one pairwise comparison for Biological Process, Cellular Component, and Molecular Function for APA genes in the 3’-most exons. Z-scores, directionality of APA changes, brain regions, and sex are represented using a combination of categorical and continuous scales, as well as different colors. **B)** Z-score normalized hierarchically clustered heatmap summarizing statistically enriched (p.adjust ≤ 0.05) GO terms in at least one pairwise comparison for Biological Process, Cellular Component, and Molecular Function for intronic APA genes. **C)** Z-score normalized RichFactor from the clusterProfiler are hierarchically clustered to create heatmaps of common cellular component GO terms for the APA genes in the 3′-most exons. **D)** Z-score normalized RichFactor from the clusterProfiler are hierarchically clustered to create heatmaps of common cellular component GO terms for the intronic APA genes. Ancestor terms (the first term immediately following the cellular component designation) are displayed by color on the left. Heatmaps are segmented based on these terms (rows). Columns are clustered based on similarity of the z-scores calculated based on the RichFactor. GO terms associated with synapses are highlighted. The number of asterisks denotes significance as follows: * ≤ 0.05; ** ≤ 0.01; *** ≤ 0.001. **E)** Venn diagram of unique APA genes in the 3′-most exon, IPA and genes that were linked to CC GO Terms containing the word stub “synap” in the description of the cellular component.

Therefore, we directed our focus toward the CC terms, as our analysis indicated that a substantial portion of our categories—based on brain region, sex, and directionality—were influenced in both REDu and REDi genes. Many of these GO terms were linked to the CC terms in shortened REDu and REDi genes, as well as lengthened genes in the prefrontal cortex of male mice.

To further identify common themes in GO terms, we identified the first ancestor term in the CC category and clustered the CC heatmap based on this information (Fig. 4C, D). This analysis revealed that a substantial part of the terms was involved in cell junction, cellular anatomical entity, intracellular anatomical structure, somatodendritic compartment, and plasma membrane, including child terms such as glutamatergic synapse, synaptic vesicle membrane, and myelin sheath for both REDu and REDi genes (Fig. 4C). CC terms related to synapses occurred in 11 of the 68 terms involved in synaptic plasticity, maintenance, and development, the majority of which were enriched in male shortened genes in all brain regions. While not all of the other CC terms reached statistical significance (p.adjust ≤ 0.05), we noticed that the terms nevertheless showed similar enrichments and were clustered together. CC terms that were significant in the lengthened genes in the male prefrontal cortex showed clustering and enrichment but to a smaller extent. BP terms associated with trans-synaptic signaling (GO:0099537), modulation of excitatory postsynaptic potential (GO:0098815), positive regulation of excitatory postsynaptic potential (GO:2000463), and excitatory postsynaptic potential (GO:0060079) were found to be significantly enriched in male hypothalamus (Table S4).

We examined the number of APA genes that overlapped across different brain regions and their directionality (Fig. 1D, E, 2B and C) with the CC GO Terms related to “synapse.” An UpSet plot visualized the unique genes found in these overlaps and the unique genes associated with the identified GO Terms (Supplementary Fig. 4B). We identified only one gene, *Map1b*, that appeared in all four categories; however, most genes did not overlap across these categories. Notably, we found that ∼42% of the REDu genes overlapped with the significant GO Terms genes related to “synapse” (Fig. 4E). Overall, enrichment of GO terms in many of our categories indicated that alcohol dependence and withdrawal affected potential structural modifications to the underlying neuronal and synaptic structures affecting preferentially the male alcohol-dependent mice.

### APA genes are associated with neurons while DE genes are associated with non-neuronal cells

To determine the association of APA and DEGs with specific cell types in the specific brain regions, we used the Allen Brain Atlas single nucleus database (*38*) to associate the APA (REDu and REDi) and DE genes with a specific cell type across the brain regions and sex (Supplementary Fig. 5, Table S6). Hierarchically clustered heatmaps based on hypergeometric distribution probabilities showed that the REDu genes were more likely to be associated with glutamatergic neurons, followed by GABAergic neurons (Fig. 5A). The region that did not show such associations was the female prefrontal cortex, which showed association of the REDu genes with microglia and pericytes. DEGs in all male comparisons were associated with astrocytes. The other associations of DEGs were with endothelial cells and microglia in various brain regions and sex as originally reported (*25*). Notably, DEGs in female hypothalamus were associated with glutamatergic and GABAergic neurons. REDi genes showed association with glutamatergic neurons across all comparisons as well as astrocytes and microglia in female amygdala.

**Figure 5.**
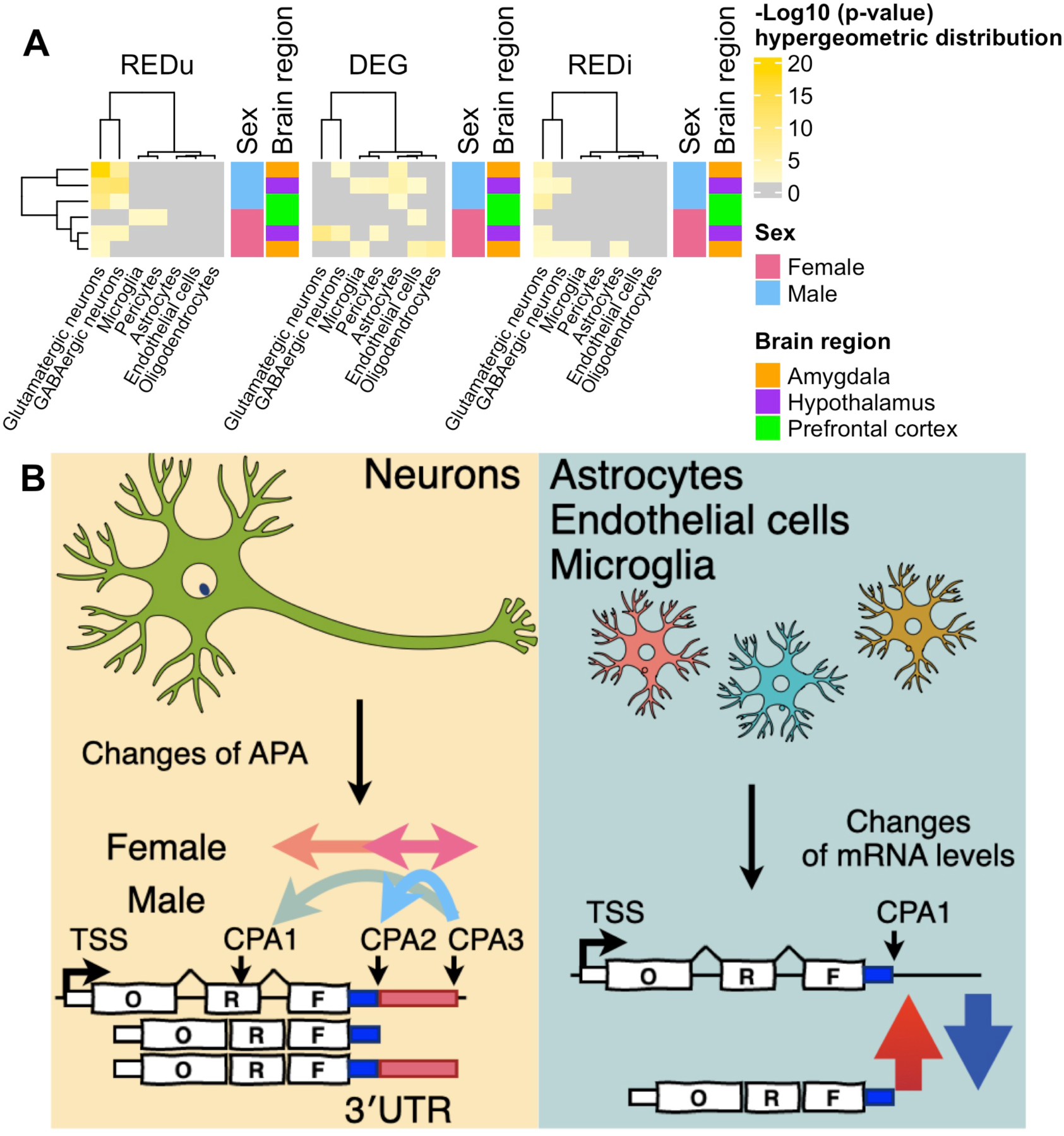
Association of APA and DE genes with distinct cell types in specific brain regions. **A)** Hierarchically clustered heatmap based on a hypergeometric distribution test, demonstrating the enrichment of REDu, DE, and REDi genes across specific cell types. - Log10 transformed p-values, brain regions, and sex are represented using a combination of categorical and continuous scales, as well as distinct color coding. **B)** Schematic representation of our model illustrating the alternative mode of regulation by alternatively polyadenylated and differentially expressed genes in alcohol-depended mice.

## DISCUSSION

In this study, we sought to determine the changes in APA in a mouse model of alcohol dependence. Our results demonstrated that alcohol dependence altered APA differently in male and female mice. In both sexes, APA genes were primarily associated with neuronal expression and specifically affecting genes in the synaptic compartment. The minimal overlap between DE and APA genes suggests that APA is an additional layer of gene expression regulation that is disrupted in AUD. While APA is a common mechanism for regulating gene expression (*3, 39*), it is frequently overlooked in neurophysiological studies since it is relatively difficult to assess with standard RNA-Seq technologies. Here, we demonstrated that alcohol dependence altered the APA of genes involved in synaptic plasticity and signaling in a sex-dependent manner across multiple brain regions, distinctions that were not apparent when enumerating differentially expressed genes (Fig. 5B). Our findings extend previous work (*22, 23, 25*) on alcohol-induced transcriptomic changes by highlighting brain region- and sex-specific APA differences, suggesting more nuanced changes of transcriptomic landscape in AUD and alcohol dependence. It is worth noting that the APA changes happened during protracted withdrawal and were likely in response to maladaptive plasticity within neural circuits that regulate reward and motivation leading to high ethanol consumption rather than the effects of acute intoxication or early withdrawal (*26*). This foundational work also linked the regulated APA genes in alcohol-dependence to neurons, suggesting that APA is an additional layer of gene expression regulation that may affect expression of specific genes in the synaptic compartment.

It has been proposed that lengthened mRNA transcripts are more likely to be associated with neurites and synapses whereas the shorter isoforms are likely to be associated with neuronal bodies (*7, 40, 41*). Although, this mechanism of mRNA localization might be more complex depending on cis-regulatory RNA localization sequences (*11, 42*), we observed what seemed to be a coordinated shortening of mRNAs encoding proteins involved in synapse strengthening and lengthening of genes involved in synapse weakening. Thus, suggesting that these perturbed isoforms might lead to modification of synaptic plasticity in AUD in a sex-dependent manner of the genes mentioned here (Fig. 4E), while explaining the maladaptive neuronal responses to increased alcohol consumption and dependence. While these findings are consistent with potential roles for APA in regulating transcripts involved in synaptic structure and signaling, further experimental studies will be required to determine whether these APA changes directly influence synaptic plasticity or neuronal function during alcohol dependence.

APA genes were most enriched in glutamatergic neurons in male mice (Fig. 5A). Functional group overrepresentation analysis also identified GO terms enriched for synaptic structural components and signaling (Fig. 4C–E, Table S4 and S5). While diverse BP, CC, and MF GO terms were uncovered by our analyses, we focused our attention on the APA changes that might affect synaptic plasticity, which could indirectly influence the long-term potentiation (LTP) and/or depression (LTD) in the brain that are associated with protracted alcohol withdrawal (*43*). Together, these mechanisms allow the brain to selectively strengthen or reduce specific neuronal circuits, shaping how information is stored and updated. By fine-tuning synaptic strength, LTP and LTD play a critical role in forming, stabilizing, and refining memories over time (*44*). Indeed, LTP was shown to be impaired during protracted withdrawal from self-administration of alcohol (*45, 46*) and our analyses identified BP terms that were related to LTP (Table S4).

Synaptic strengthening is mediated by the activation of NMDA receptors, resulting in calcium influx and subsequent activation of CaMKII, a central organizer of synaptic plasticity, learning and memory (*47, 48*). CaMKII activation enhances the formation of scaffolds at the postsynaptic side through the accumulation of PSD proteins and actin polymerization. In contrast, the decrease in synaptic strength is characterized by lower calcium influx through NMDA receptors, activating calcineurin, which trigger the internalization of AMPA receptors and weakens synaptic transmission. Our analyses identified fourteen genes within the CaMKII family (Table S2). Of these, six members contained only a single CPA site and were unlikely to be regulated on APA level, while the remaining eight had multiple CPA sites. Two genes (*Camk2a*, *Camk2b*) were shortened in the 3′-most exon across all male brain regions and one gene (*Camk2n1*) demonstrated shortening in all comparisons (Table S2). Another gene, *Camk1d,* was shortened in all male brain regions but as an REDi gene. Similarly, NMDA receptor genes that possess a single CPA site do not appear to be affected by APA (Table S2). AMPA receptor genes had no detectable APA changes, although multiple CPA sites were identified in each (Table S2).

Other genes involved in synaptic plasticity were affected by APA in male alcohol-dependent mice. We identified shortening of the *Olfm1* gene (aka *Noelin*), which modulates lateral mobility of AMPA receptors and thus is involved in synapse strengthening (*49*). *Map1b,* which is required for the internalization of AMAPR and is important in synapse weakening (*50*), was also shortened. Conversely, we determined that *Ppp3ca*, a gene that encodes the alpha isoform of a subunit of calcineurin (*51*), a powerful inhibitor of synapse strengthening, was lengthened in its 3′-most exon.

We did not find an enrichment of the APA genes in oligodendrocytes or oligodendrocyte precursor cells (Fig. 5A). However, one enriched GO term linked to APA changes was myelin sheath (Fig. 4C and Table S4). Closer examination of these genes (Table S4) indicated that their expression was mostly associated with neurons (Table S6). Indeed, recent work has suggested that neuronal activity can modulate adaptive myelination specifically in memory formation and learning (*52*). Formation of myelin has been reported to be reduced in AUD (*53, 54*). Likely, myelin sheath reduction might be dynamically remodeled in response to impaired neuronal synapse strength associate with alcohol dependence and linked to APA genes in the neuronal compartment.

In male mice, we found that hundreds of APA genes across all three brain regions exhibited shortening, while female mice did not demonstrate a clear pattern of change. In females, the number of APA genes varied by brain region and was nearly ten times fewer than those in the corresponding male brain regions. We were unable to provide a solid rationale for the lower number of APA genes and the lack of an apparent pattern in females. However, we observed that, generally the female mice had shorter mRNA transcripts compared with the males. Given the current datasets, we could not disentangle the contributions of sex and increased ethanol consumption to what seemed to be an even greater sex-dependent shortening of APA in alcohol-dependent females compared with males.

Elevated alcohol consumption leads to activation of the innate immune system. Proinflammatory cytokines are increased in the blood and brains of humans with AUD (*55*). Analysis of the REDi genes indicated that again in males there was a pattern of shortening. A substantial number of the CPA sites of these 34 shortening genes in all three male brain regions were in internal exons (Table S3). Accumulation of RNAs without protein coding potential but with increased propensity of forming stable secondary structures has been linked to increased neuroinflammation (*56*). High-throughput sequencing from the frontal cortex of human subjects determined that differentially expressed genes were involved in neuroinflammation, most often associated with astrocytes, oligodendrocytes, and microglia (*57-59*). Consistent with those findings, the innate immune system was activated in mouse models, which were developed based on a selective breeding of mice consuming high and prolonged ethanol exposure (*28, 60*). Indeed, systemic activators of the innate immune system such as lipopolysaccharide (LPS) and polyinosinic:polycytidylic acid [poly(I:C)] led to increased alcohol intake (*61, 62*). However, REDi genes were associated with neurons (Fig. 5A) in our studies. Therefore, the contribution of the REDi genes to the pool of double-stranded RNA in AUD and neuroinflammation is currently unclear. Also unclear is whether the selection of CPA sites in internal exons is due to a specific molecular mechanism activated by protracted alcohol withdrawal or is a stochastic event. Further studies will be necessary to answer these questions.

Overall, we propose that alternative polyadenylation is a distinct layer of gene expression regulation independent from mRNA abundance in AUD (Fig. 5B) and specifically in the synaptic compartment. Based on correlative cell type-specific marker genes identified from snRNA-Seq analysis, APA genes were primarily associated with neurons. We also showed that APA changes associated with chronic alcohol exposure were more prominent in male than in female mice. The underlying biological mechanisms for this sex-dependent effects have yet to be fully investigated. Further research will address whether alcohol dependence leads to the emergence of new CPA sites in this mouse model and in other models, including humans and their physiological relevance to AUD. Additional future studies will determine the causal relationship between APA and AUD.

## MATERIALS AND METHODS

### Datasets

109 fastq TagSeq files from specific brain regions, sex, and treatment (accession number GSE176122) (*25*) were downloaded programmatically from Gene Expression Omnibus using SRA-Toolkit. The batch 1 fastq “on-site” reads are available from GSE152975, wild type mice (SRR12067714, SRR12067715, SRR12067719) (*63*). The batch2 fastq “on-site” files are available from GSE310389 (this manuscript). Single nucleus (sn) RNA-Seq (*38*) datasets were downloaded from GitHub (https://github.com/AllenInstitute/abc_atlas_access/blob/main/descriptions/WMB-10X.md).

### Adapting the PolyA_DB version 3.2 to mm10 reference genome

Mouse PolyA_DB version 3.2, originally compiled on the mouse mm9 reference genome, was downloaded from https://exon.apps.wistar.org/polya_db/v3/ (*35*). Unique names were assigned to each CPA site, and the resulting database was processed through the liftover online tool provided by the Genome Browser to convert the mm9 CPA sites to mm10 (*64*). This process identified a total of 398,869 mm10 CPA sites. To further cross-reference the identities of these CPA sites, we utilized PolyA_DB for mm10 (mouse.PAS.mm10.rds) prepared by the authors of the MAAPER bioinformatics tool (*33*), which contained 236,455 CPA sites. The MAAPER database did not include intergenic sites; therefore, we removed the intergenic sites from the PolyA_DB mm10 database, resulting in a total of 242,674 CPA sites. After comparison, we identified a total of 228,517 unique CPA sites. Of these, 627 CPA sites were unique to the mm10 database prepared by liftover, while 390 were unique to the MAAPER mm10 database (Supplementary Fig. 1A). Additionally, 227,500 CPA sites were present in both databases. The major discrepancy in the CPA sites was observed on chromosome 10, where a total of 703 out of 1,017 sites did not overlap. This discrepancy appeared to be related to an inversion in the region on chromosome 10 from ∼93,320,570 to 100,718,892 between the mm9 and mm10 reference genomes.

### Identification of moderate to high expressing CPA sites in whole mouse brain

Fastq files obtained with “on-site” RNA sequencing technology were directly aligned on a mm10 (GENCODE GRCm38) indexed genome, which was generated using gencode.vM23.primary_assembly.annotation gtf file (created September 2019, GENCODE) using STAR v.2.7.10a_alpha_220207 (*65, 66*). For this alignment, we used the following STAR parameters --outFilterType BySJout --outFilterMultimapNmax 20 -- alignSJoverhangMin 8 --alignSJDBoverhangMin 1 --outFilterMismatchNmax 999 -- outFilterMismatchNoverLmax 0.1 --alignIntronMin 20 --alignIntronMax 1000000 -- alignMatesGapMax 1000000, and with sam attributes NH, HI, NM, MD (-- outSAMattributes). An index file was created for each Binary Alignment Map (BAM) file, and the BAM files were further processed using a modified python script based on step5.LAP.hunter.qcREV (https://github.com/RJWANGbioinfo/APAlyzer-QSr/tree/master/scripts). The counts for each BAM file were processed to identify CPA sites that were expressed by at least three reads in at least three sample using a modified merged version of step6_1.LAP2PAS.R and step6_2.Quant_R.pas2gene.builder.R, which were available from the same GitHub repository (*35, 67*).

To determine the overall similarity of the aligned reads and CPA sites, we performed a correlation analysis across the mapped reads of these samples. Similarity correlation pairwise analysis between the BAM files was performed by using multiBamSummary (deeptools v 3.5.6) (*68*) with default parameters. Spearman correlation was calculated and plotted using plotCorrelation (deeptools v 3.5.6). In all pairwise comparisons, the Spearman coefficient was larger than 0.80 (Supplementary Fig. 1B), indicating good concurrence across the brain samples and the reliability of the curated CPA sites. The final curated database (for details, see Materials and Methods) contained 44,770 CPA sites located in 15,906 genes and non-coding (nc) RNAs (Table S1). The vast majority of CPA sites were located in the 3′-most exon genomic feature, followed by introns (Supplementary Fig. 2A).

### Quality control and alignment of TagSeq reads

Quality control and trimming of the reads for Taq-Seq to 75 bp were performed using fastp v.0.23.2 tool with options -t 25, -w 6 and an HTML report was generated (*69*). On average, between 4.8 and 6.6 million reads were mapped per sex (female, male). Trimmed reads from each individual mouse and brain region were aligned on a mm10 (GENCODE GRCm38) indexed genome, which was generated using the gencode.vM23.primary_assembly.annotation gtf file (created September 2019, GENCODE) using STAR v.2.7.10a_alpha_220207 (*65, cc*). For the alignment, we used the following STAR parameters --outFilterType BySJout --outFilterMultimapNmax 20 --alignSJoverhangMin 8 -- alignSJDBoverhangMin 1 --outFilterMismatchNmax 999 --outFilterMismatchNoverLmax 0.1 --alignIntronMin 20 --alignIntronMax 1000000 --alignMatesGapMax 1000000, and with sam attributes NH, HI, NM, MD (--outSAMattributes). On average between 75% and 78% of the reads were uniquely mapped to the mouse genome.

For each sorted BAM file from TagSeq corresponding to an individual mouse, an index file was created using samtools v.1.15.1 (*70*). BAM files for each individual treatment group were merged using samtools v.1.15.1 and index file was created as explained above. Bigwig files corresponding to both strands were created using deeptools v 3.5.1 with parameters bamCoverage --binSize 1 and normalized using the CPM method (*68*). Stranded bigwig files were created by using the above parameters and either –filterRNAstrand forward or -- filterRNAstrand reverse for the forward or reverse strands including -- effectiveGenomeSize = 2652783500, respectively.

### Creation of metagene plots

As a first step, we created metagene plots covering the regions around the CPA site (-600 to +300 nucleotides) to determine the relative normalized distribution of the reads in TagSeq vs the “on-site” reads. Metagene plots were constructed using metagene2 (*71*). A metadata was used to identify CPA type, and genomic feature (intron - exon location) (Table S1). After creation of the new metagene object a metagene plot was produced with a produce_metagene with a confidence interval parameter (alpha) set to 0.01. Subsequently, the mean coverage for each bin, CPA type and genomic feature was extracted from the metagene object and further normalized using min-max approach where for each value the minimum was subtracted and divided by the difference between maximum and minimum value, thus the distribution values ranged from 0 to 1.

We separated the metagene plots by genomic features and CPA types as detailed in the polyA database (Supplementary Fig. 1C) (31). Notably, the reads located in 3′ UTR (F), 3′ UTR (M), 3′ UTR (L), 3′ UTR (S), and introns were consistent and uniform for the “near-site” and “on-site” reads with well-defined peaks and distances between the peaks around the CPA sites and a narrow confidence interval. Whereas the reads in CDS, ncRNAs, pseudogenes, and 5′ UTR formed peaks that were more broadly distributed around the CPA sites and did not exhibit well-defined boundaries for either “near-site” or “on-site” reads with a larger variation in the coverage. Furthermore, the normalized density coverage of the reads relative to the CPA in the training genes was uniform, with a peak of ∼ 260 nucleotides away from the CPA site across the samples, suggesting that the training models were consistent across brain region and sex (Supplementary Fig. 2C). As an additional quality control measure for TaqSeq across the samples, we generated normalized coverage metagene plots, which were segmented by CPA type (3′UTR (F), 3′UTR (M), 3′UTR (L), 3′UTR (S), and Intron), brain region, and sex (Supplementary Fig. 2D). The distribution and the peaks for these metagene plots were also consistent across brain region and sex and aligned well with the normalized density distribution of the training genes (Supplementary Fig. 2C).

Next, we used the MAAPER software (32) to determine APA events. MAAPER predicts the distribution of the alignment hits without assuming data uniformness from “training” genes possessing only a single CPA site (31, 32). Our analysis revealed that all samples exhibited a similar number of “training” genes that did not differ statistically across the pairwise comparisons between alcohol- and non-dependent mice (with a mean of 1,907 genes, range 1,120 to 2,550, Supplementary Fig. 2B), resulted in a comparable distribution of reads relative to the single CPA sites (Supplementary Fig. 2D).

### MAAPER analysis and additional custom functions

To facilitate the processing of multiple brain regions in male and female mice, we utilized several custom functions to automate the analyses. A final object corresponding to the gene file from the MAAPER containing all the pairwise comparison was created (Table S2). We assigned our curated polyA database (see above) to the MAAPER’s parameter pas_annotation. We assigned the same gtf file that was used in the BAM alignment to MAAPER’s parameter gtf. The length parameter (read_len) was set to 75.

MAAPER uses the likelihood ratio test (LRT) and Fisher’s exact (FET) tests to determine whether the gene’s APA profile differs between alcohol- and non-dependent mice. A FET was applied to the two CPA sites with the greatest change in ratio to determine whether the use of these CPA sites differed statistically across the pairwise comparisons. To monitor the false positive rate resulting from multiple comparisons, we applied Benjamini-Hochberg’s (BH) test (*72*) using p.adjust() and calculated adjusted p-values for LTR and FET independently (Table S2).

We used several R packages [ComplexUpset plots (*34*), ComplexHeatmaps (*73*) and eulerr (*73*)] to produce the UpSet plots, heatmaps, and proportional venn/eluer diagrams. To keep sample alignment consistent, we first performed unsupervised hierarchical clustering on the REDu category to generate a column dendrogram. This clustering order was then applied to the DEG and REDi heatmaps. For Figure 5A, the row dendrogram was similarly derived and applied across all categories. Additional custom functions in R were produced and implemented to adapt the analyses and visualize the results.

### Functional group overrepresentation analysis

We used the clusterProfiler package in R (*74*) to determine GO terms that were enriched based on the APA genes. Statistically significant APA genes that were shortened and lengthened were used independently to identify enriched GO terms. The “universe” (background) genes were set to all unique gene symbols in all six comparison that did not have NA value in the APA analysis for both REDu and REDi. p-value and q-value for the analysis were set to 1 and the multiple test comparison correction method was set to “BH”. Further we used the “simplify” function with “by” set to “p.adjust”, “cutoff” to 0.7, “select_fun” to “min” and the “measure” to “Wang.” All gene ontology terms (Biological Process, Cellular Component, and Molecular Function) were queried in a sequential approach. Further we have selected the terms that show statistical significance (p.adjust ≤ 0.05) in any of the six comparison and extracted these terms regardless of their significance status in the rest of the comparisons. We used the z-score normalized RichFactor to construct the heatmaps. To summarize the number of statistically significant events (p.adjust ≤ 0.05) GO terms per comparison, we constructed additional z-score corrected heatmaps. Ancestor terms were extracted and the heatmaps were split based on these ancestor terms. Two GO terms, one for REDu and REDi genes did not have a specific ancestor term and were manually assigned to cell-junction and protein-to-protein complex. Further we extracted the GO terms having “synap” anywhere in their GO term Description and linked them to the heatmaps. Additionally, the GO Term myelin sheath was identified to be the only CC term with a p.adjusted ≤ 0.01 in four comparisons (Figure 4). Therefore, myelin sheath was manually added to the displayed terms.

### Cell type overrepresentation analysis using the Allen brain atlas datasets

We downloaded all 24 dataset files and their associated metadata from the GitHub repository (see above) (*38*) and consolidated the single nucleus matrix data into a single list object in R. Subsequently, we extracted cells from the prefrontal cortex (identified by the “region_of_interest_acronym” matching the “PL-ILA-ORB” descriptor), the amygdala (identified by the “region_of_interest_acronym” matching “CTXsp”), and the hypothalamus (identified by “feature_matrix_label” matching “-HY”). Following an additional filtering step (see our deposited scripts), we relabeled cells with a “cluster_annotation_term_name” equal to “Glut” as glutamatergic neurons, cells labeled as “GABA” as GABAergic neurons, and those labeled as “Astro” as astrocytes. Cells identified as “Oligo-OPC” were categorized as oligodendrocytes. Endothelial cells were designated based on the cluster_alias column with values of 5252, 5254, and 5255; pericytes were identified by cluster_alias values of 5243 and 5248; microglia by cluster_alias value of 5283; and smooth muscle cells by cluster_alias values of 5244 and 5249. Any remaining cells that did not receive a specific label based on these criteria were removed from the dataset. These labels were added to the meta.data in the respective Seurat objects (*75*) and used further to calculate the mean expression of the genes per cell types (Table S6) and to perform the hypergeometric distribution analyses. We used an arbitrary value equal or bigger than 0.4 in the mean expression table to associate a gene with cell type-specific expression. To determine the probability of specific genes being associated with specific cell types, we used hypergeometric distribution function phyper() with “lower.tail” set to “FALSE.” Heatmaps were built with ComplexHeatmaps. Initially, the columns (representing the cell types) and rows (representing the brain region and sex) were allowed to cluster based on the APA genes in the 3′-most exon (REDu). The specific column and row dendrograms were applied to the other categories. Negative Log10 transformed p-values were displayed in shades of yellow in the corresponding cells if they were bigger than 1.3.

## ACKNOWLEDGEMENTS

We thank Christian Bustamante for advice on the use of R studio and other bioinformatical tools. During the preparation of this work, the authors used AI-powered language processing models, as a language assistant in order to improve written text and to optimize the code used in the analyses. After using these tools/services, the authors reviewed and edited the content as needed and take full responsibility for the content of the publication.

## Author contributions

Conceptualization, P.N.G.; Methodology, P.N.G., C.C.M. and I.P.; Formal analysis, P.N.G.; Writing—Original Draft, P.N.G.; Writing—Review and Editing, P.N.G., B.R.K, C.C.M., L.B.F., R.O.M., and I.P.; Supervision, P.N.G.

## Conffict of interest statement

None declared.

