## Supplementary Figures for "Alternative polyadenylation in the brain is altered by chronic ethanol exposure in a sex- and cell type-specific manner"

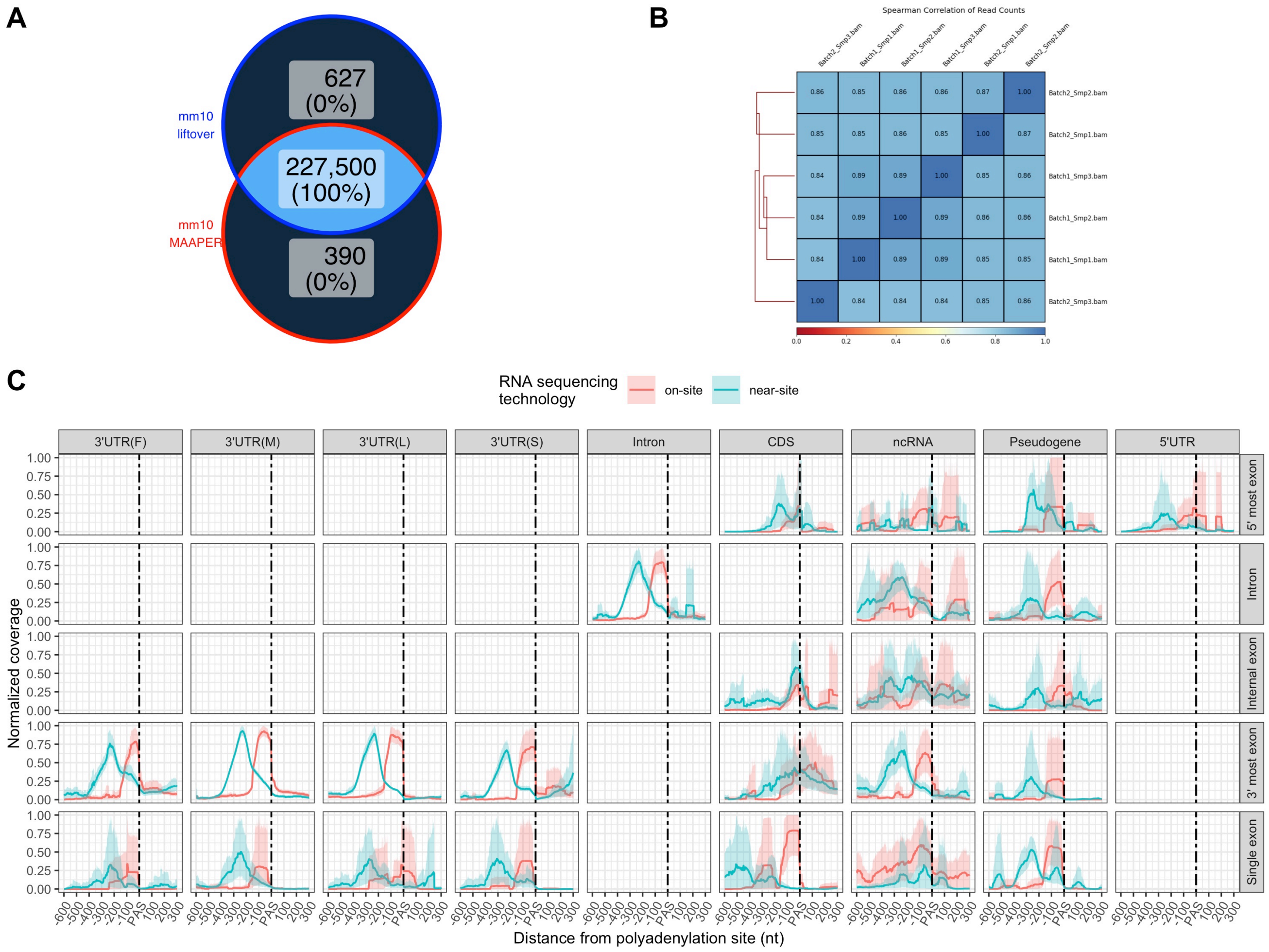
A
mm10
liftover
mm10
MAAPER
627
(0%)
227,500
(100%)
390
(0%)
B
Spearman Correlation of Read Counts
Batch2_Smp3.bam
Batch1_Smp1.bam
Batch1_Smp2.bam
Batch1_Smp3.bam
Batch2_Smp1.bam
Batch2_Smp2.bam
0.86 0.85 0.86 0.86 0.87 1.00 Batch2_Smp2.bam
0.85 0.85 0.86 0.85 1.00 0.87 Batch2_Smp1.bam
0.84 0.89 0.89 1.00 0.85 0.86 Batch1_Smp3.bam
0.84 0.89 1.00 0.89 0.86 0.86 Batch1_Smp2.bam
0.84 1.00 0.89 0.89 0.85 0.85 Batch1_Smp1.bam
1.00 0.84 0.84 0.84 0.85 0.86 Batch2_Smp3.bam
0.0 0.2 0.4 0.6 0.8 1.0
C
RNA sequencing technology
on-site
near-site
3'UTR(F)
3'UTR(M)
3'UTR(L)
3'UTR(S)
Intron
CDS
ncRNA
Pseudogene
5'UTR
5' most exon
Intron
Internal exon
3' most exon
Single exon
Normalized coverage
0.00 0.25 0.50 0.75 1.00
-600 -500 -400 -300 -200 -100 PAS 100 200 300
Distance from polyadenylation site (nt)

**Supplementary Figure 1. Curated PolyA database. A)** Venn diagram illustrating the CPA sites identified through the liftover method, alongside those available via MAAPER for the mm10 reference genome. **B)** Heatmap showing pairwise Spearman correlation coefficients for the “on-site” BAM files, utilized to identify moderate to high expressing transcripts and their corresponding CPA sites. **C)** Metagene plots for “on-site” and “near-site” sequencing technologies, segmented according to genomic features and CPA types as defined in PolyA_DB.

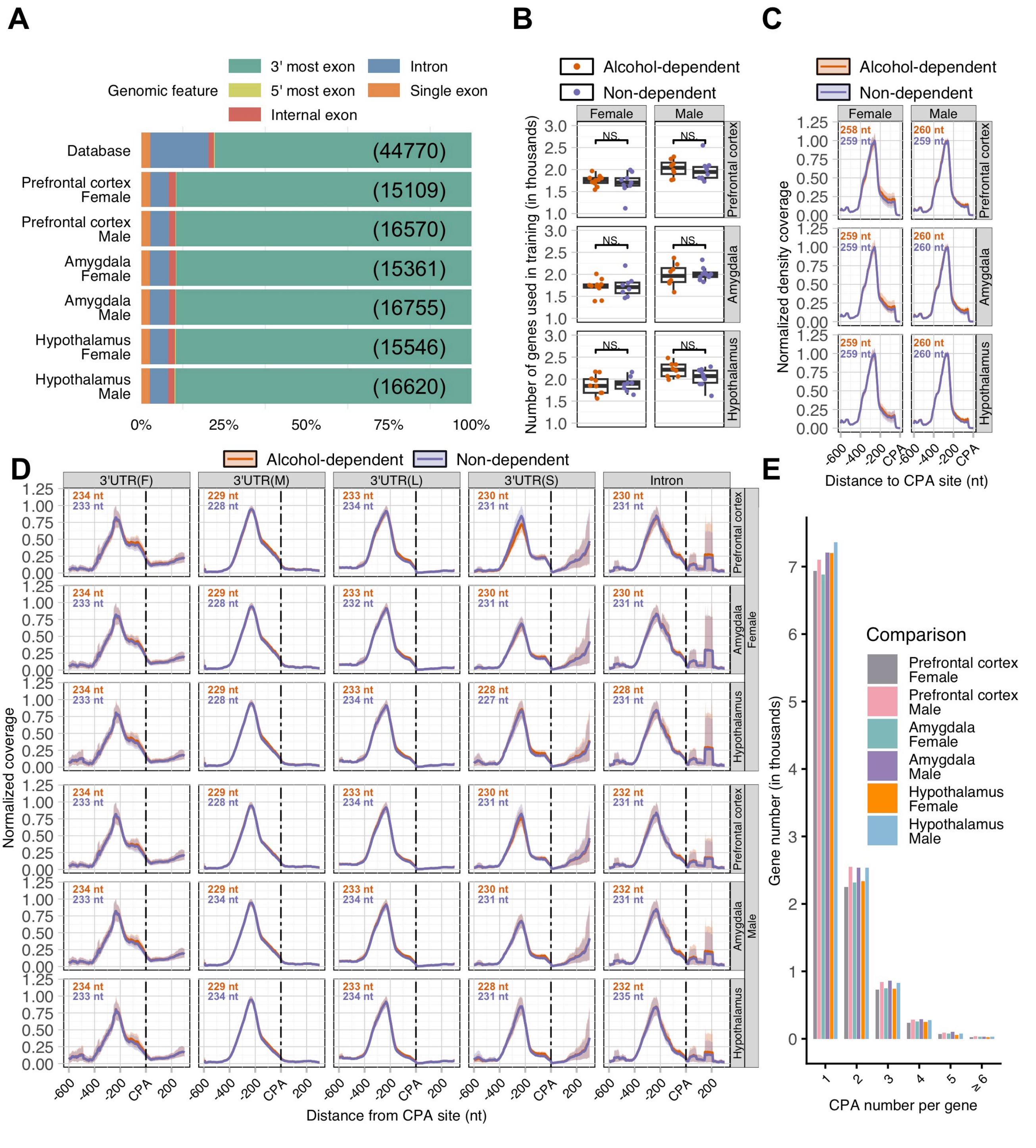
A
Genomic feature
3' most exon
5' most exon
Internal exon
Intron
Single exon
Database
(44770)
Prefrontal cortex Female
(15109)
Prefrontal cortex Male
(16570)
Amygdala Female
(15361)
Amygdala Male
(16755)
Hypothalamus Female
(15546)
Hypothalamus Male
(16620)
0%
25%
50%
75%
100%
B
Alcohol-dependent
Non-dependent
Female
Male
Prefrontal cortex
Amygdala
Hypothalamus
NS
Number of genes used in training (in thousands)
C
Alcohol-dependent
Non-dependent
Female
Male
258 nt
259 nt
260 nt
259 nt
260 nt
Normalized density coverage
Distance to CPA site (nt)
D
Alcohol-dependent
Non-dependent
3'UTR(F)
3'UTR(M)
3'UTR(L)
3'UTR(S)
Intron
Prefrontal cortex
Amygdala
Hypothalamus
Female
Male
Normalized coverage
Distance from CPA site (nt)
E
Comparison
Prefrontal cortex Female
Prefrontal cortex Male
Amygdala Female
Amygdala Male
Hypothalamus Female
Hypothalamus Male
Gene number (in thousands)
CPA number per gene
≥ 6

**Supplementary Figure 2. TagSeq as a reliable method for APA analysis. A)** Total number and percent distribution of CPA sites across genomic features for the database, along with pairwise comparisons between animals subjected to chronic intermittent ethanol (CIE) exposure (alcohol-dependent) and control mice exposed to air (non-dependent), separated by sex and brain region. **B)** Count of single CPA genes included in the MAAPER model training, referred to as training genes. No statistical difference was observed in the pairwise comparisons, as indicated by “NS” for alcohol versus non-dependent mice. **C)** Normalized density coverage of read distribution for the “training genes” in pairwise comparisons within a 600-nucleotide region upstream of the CPA sites in alcohol-dependent versus non-dependent mice. **D)** Metagene plot illustrating the normalized coverage distribution of reads across CPA types as derived from PolyA_DB (*35*), segmented by sex and brain region for alcohol- and non-dependent mice. **E)** Bar graph displaying the number of genes with a specific count of CPA sites in the pairwise comparisons.

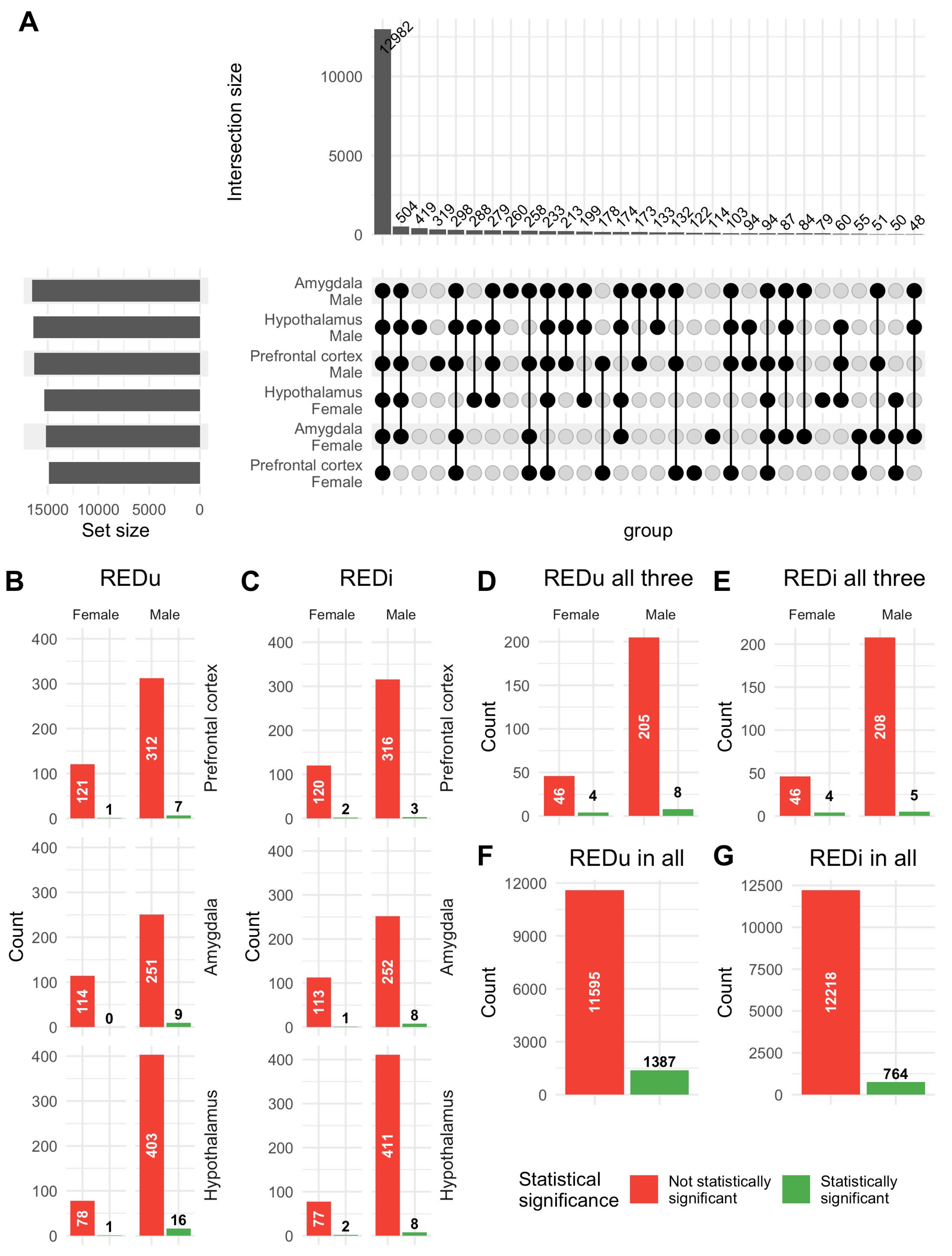
A
Intersection size
12982
504 419 319 298 288 279 260 258 233 213 199 178 174 173 133 132 122 114 103 94 94 87 84 79 60 55 51 50 48
Amygdala Male
Hypothalamus Male
Prefrontal cortex Male
Hypothalamus Female
Amygdala Female
Prefrontal cortex Female
Set size
group
B REDu
C REDi
D REDu all three
E REDi all three
F REDu in all
G REDi in all
Female
Male
Count
Prefrontal cortex
Amygdala
Hypothalamus
121 1 312 7
114 0 251 9
78 1 403 16
120 2 316 3
113 1 252 8
77 2 411 8
46 4 205 8
46 4 208 5
11595 1387
12218 764
Statistical significance
Not statistically significant
Statistically significant

**Supplementary Figure 3. Overlapping and Unique CPA Sites in All Pairwise Comparisons. A)** UpSet plots illustrating all CPA sites across pairwise comparisons. Bar graphs depict the unique CPA sites categorized by brain region and sex, alongside the number of these sites identified as being linked to statistically significant events in REDu (**B**) and REDi (**C**) genes. Additionally, bar graphs display the overlapping CPA sites identified exclusively in the brain regions of female and male mice, along with the count of these sites associated with statistically significant events in REDu (**D**) and REDi (**E**) genes. Panels **F)** and **G)** represent the distribution of statistically significant events within the CPA sites that overlapped across all six comparisons for REDu and REDi.

A

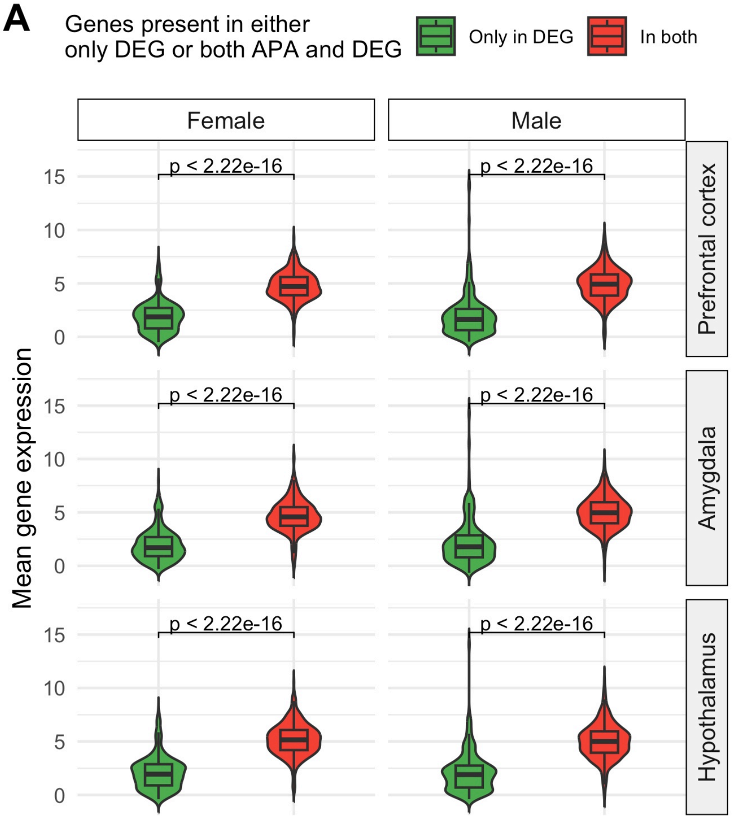

Genes present in either only DEG or both APA and DEG
Only in DEG
In both
Female
Male
Prefrontal cortex
Amygdala
Hypothalamus
Mean gene expression
p < 2.22e-16


B

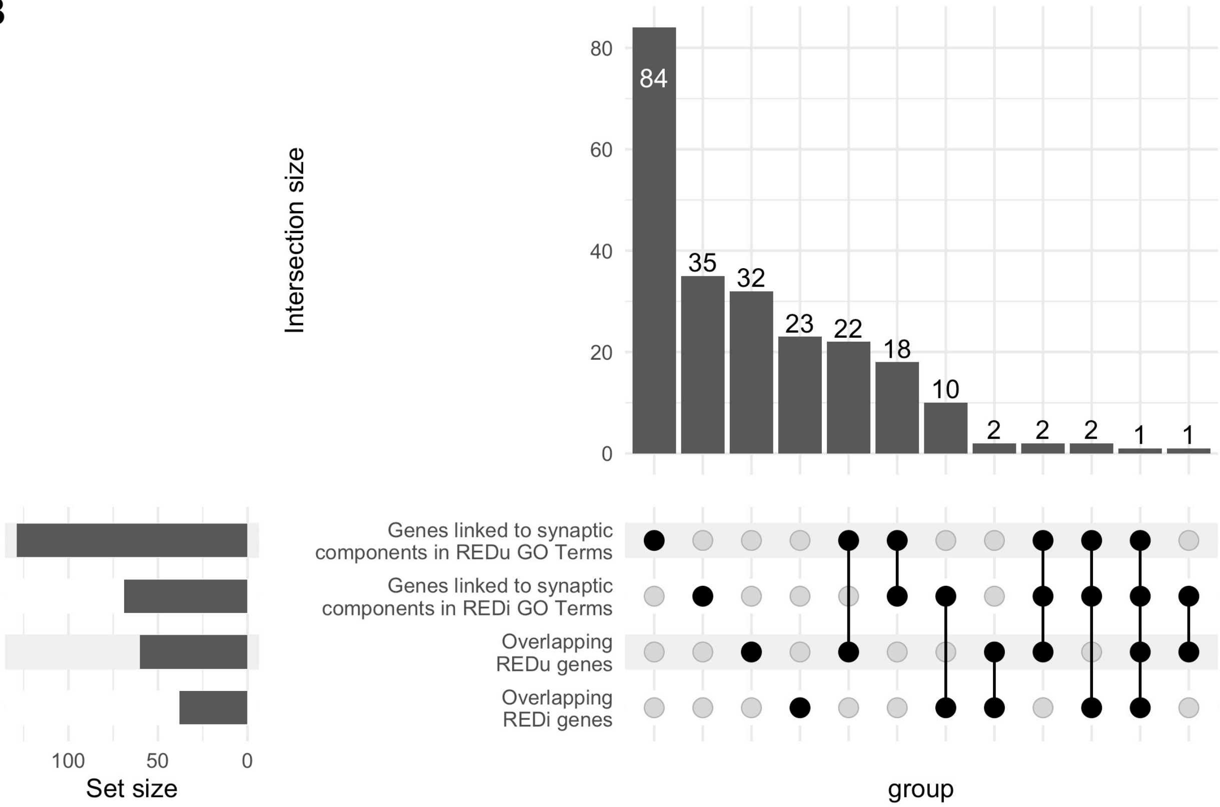

Intersection size
84
35
32
23
22
18
10
2
2
2
1
1
Genes linked to synaptic components in REDu GO Terms
Genes linked to synaptic components in REDi GO Terms
Overlapping REDu genes
Overlapping REDi genes
Set size
group

**Supplementary Figure 4. A)** Pairwise comparison of the mean expression of genes that were not mapped to the MAAPER gene output (only in DEG, shown in green) versus those mapped to both (depicted in red). Statistical significance, defined as a $p < 0.05$, is indicated by a Wilcoxon test. **B)** UpSet plot of unique APA genes in the 3'-most exon, intron and genes that were linked to CC GO Terms containing the word stub “synap” in the description of the cellular component for both APA categories.

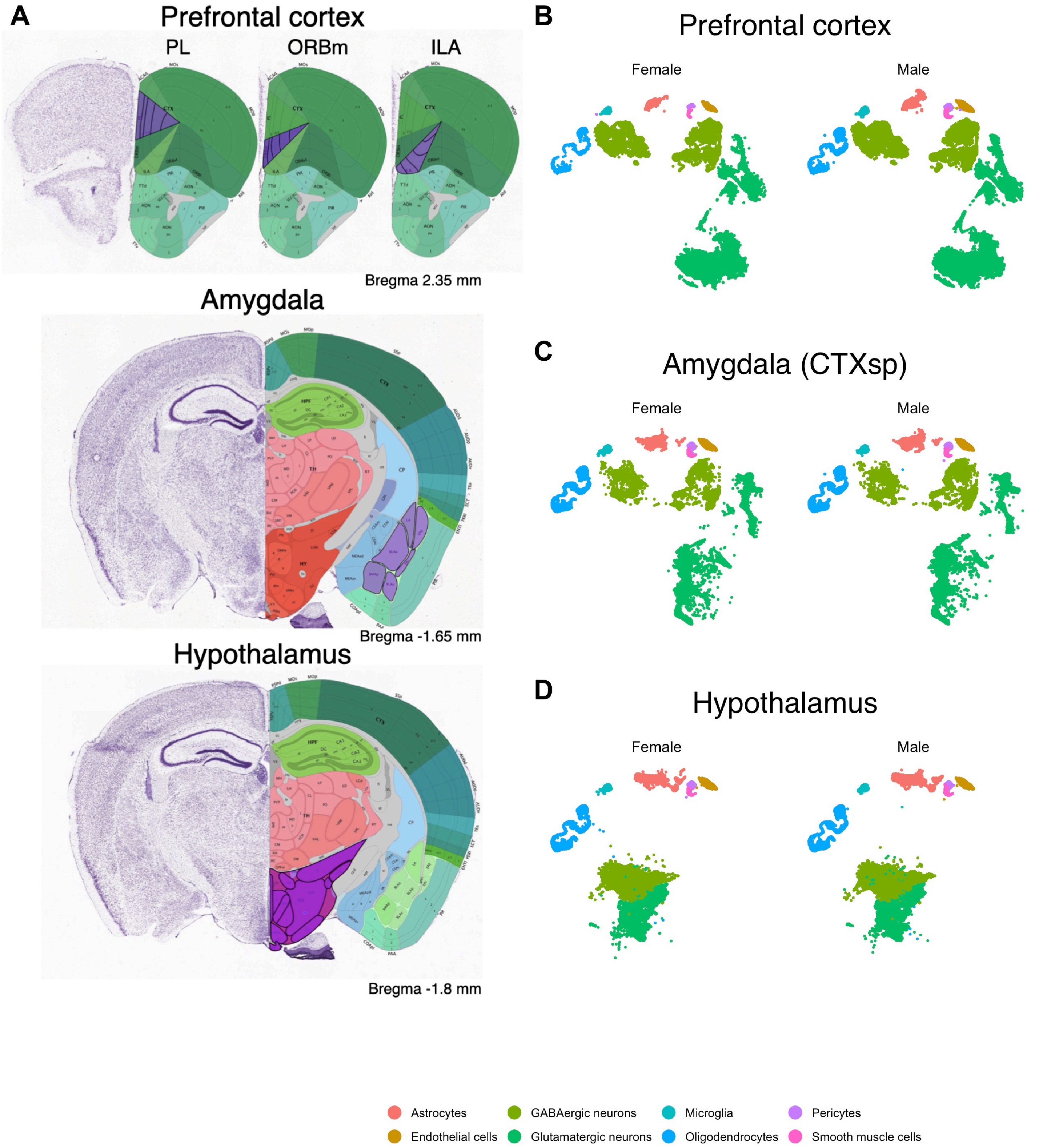
A
Prefrontal cortex
PL
ORBm
ILA
Bregma 2.35 mm
Amygdala
Bregma -1.65 mm
Hypothalamus
Bregma -1.8 mm
B
Prefrontal cortex
Female
Male
C
Amygdala (CTXsp)
Female
Male
D
Hypothalamus
Female
Male
Astrocytes
GABAergic neurons
Microglia
Pericytes
Endothelial cells
Glutamatergic neurons
Oligodendrocytes
Smooth muscle cells

**Supplementary Figure 5. A)** Coronal sections of the mouse brain, derived from the Allen Brain Atlas (https://mouse.brain-map.org/experiment/thumbnails/100142143 and mouse.brain-map.org), illustrating the prefrontal cortex, amygdala, and hypothalamus that were sampled in the original study (*25*). The mean bregma coordinates from the original study are indicated in the lower right corner of the respective brain regions. Areas of interest are highlighted in violet. **B)** UMAPs depicting single-nucleus data for the specified brain regions were generated based on the x and y coordinates from the original study (*38*), further segmented by sex and cell types.
